# Genesis of Ecto-symbiotic features based on Commensalistic Syntrophy

**DOI:** 10.1101/2022.09.04.506536

**Authors:** Nandakishor Krishnan, Villő Csiszár, Tamás F. Móri, József Garay

**Affiliations:** Institute of Evolution, Centre for Ecological Research, Konkoly-Thege M. út, H-1121, Budapest, Hungary; Doctoral School of Biology, Institute of Biology, Eötvös Loránd University, Pázmány Péter s. 1/C, H-1117, Budapest, Hungary; Department of Probability Theory and Statistics, Eötvös Loránd University, Pázmány Péter s. 1/C, H-1117, Budapest, Hungary; Alfréd Rényi Institute of Mathematics, Reáltanoda u. 13-15, H-1085, Budapest, Hungary

**Keywords:** Ecto-symbiosis, Commensalism, Syntrophy, Evolutionary substitution, Coexistence

## Abstract

The origin of eukaryotes and organellogenesis have been recognized as a major evolutionary transition and subject to in-depth studies. Acknowledging the fact that the initial interactions and conditions of cooperative behaviour between free-living single-celled organisms are widely debated, we narrow our scope to a single mechanism that could possibly have set-off multi-species associations. We hypothesize that the very first step in the evolution of such cooperative behaviour could be a single mutation in an ancestral symbiont genome that results in the formation of an ecto-commensalism with its obligate ancestral host. We investigate the ecological and evolutionary stability of inter-species microbial interactions with vertical transmissions as an association based on syntrophy (cross-feeding). To the best of our knowledge, this is the first time that a commensalistic model based on the syntrophy hypothesis is considered in the framework of coevolutionary dynamics and invadability by mutant phenotype into a monomorphic resident system.

## Introduction

Nature has devised mechanisms for some species to coexist in associations in which organisms of one species are biologically linked to organisms of another species, and examples of these interactions are frequently observed and extensively studied. Though definitions vary, symbiosis is often described as a long-term and intimate relationship of individuals of different genome in which partners may or may not live in physical contact (often physiologically integrated) and have already started to coevolve. However, there is ambiguity in the definition of symbiosis and the kinds of species interactions that should fall under its purview (Martin and Schwab^1^, 2013). In this paper, we adopt the above conventional definition of “symbiosis” (Paracer and Ahmadjian^2^, 2000) which could be obligate (Kaur et al.^3^, 2021; Moya et al.^4^, 2008). The partnership can be mutually beneficial (mutualism) or can be asymmetrical with or without a conflict of interest (parasitism or commensalism). Ecological interactions are often complicated and can be placed on a continuum varying from fitness-reducing parasitism to fitness-increasing mutualism (Ewald^5^, 1987; Smith and Szathmary^6^, 1997), and these host-symbiont relationships have a long history of coevolution (Kaur et al.^3^, 2021; Moya et al.^4^, 2008).

The long-term biological interactions between two different organisms can occur as a consequence of horizontal and/or vertical transmission. The subsequent evolution of associations that probably started as a commensal one has been towards parasitism in species with horizontal transmission while vertical transmission promotes mutualism (Herre^7^, 1993; Bright and Bulgheresi^8^, 2010). Evolution from horizontal transmission and parasitism towards vertical transmission and mutualism has also been discussed (Clay^9^, 1988; Afkhami and Rudgers^10^, 2008). Thus, there are evidence to support the argument that evolution towards mutualism or parasitism is highly influenced by the mode of transmission. The vertical transmission (when the symbionts are directly transferred from an infected host to its offspring) guarantees the association of host and its symbiont in the immediate generation. One of the main consequences of vertical transmission is that the evolutionary success of the symbiont and its host is determined by that of the host-symbiont association (Camus et al.^11^, 2022). In other words, the vertical transmission opens the door to the coevolution of host-symbiont pair in the direction of mutualism. Thus, the emergence of vertical transmission should be one of the important questions in the evolution of symbiosis and is one of the focuses of our study.

We develop an ecological model that describes interactions leading to symbiosis of unicellular and free-living (physically independent) individuals, and general conditions that eventually facilitate and stabilize such interactions. Here we are interested in the first step in the evolutionary origin of a vertically transmitted obligate symbiosis, i.e., the evolution of coexistence between two free-living species (Nguyen and van Baalen^12^, 2020). In other words, the resident ecological system was formed by two physiologically independent species, and existence and survival of at least one of these species is dependent on the other (commensalistic). This ecological dependence between these species is crucial for theoretical modelling perspectives too. The model can be extended to understand potential endosymbiosis, especially of those originated from mitochondrial integration. For instance, if the host ancestor predated the symbiont ancestor by phagocytosis, that could have eventually opened the door for endosymbiosis (when the symbiont ancestor could survive within the host) (Zachar et al.^13^, 2018). Clearly, in this scenario the vertical transmission immediately occurs since the symbiont lives within the host ancestor cell and thereby endosymbiosis is evolved. The host-symbiont coexistence is thereby an important issue in the evolution of the monophyletic eukaryotes (Lake^14^, 1987). According to the endosymbiotic theory (Margulis^15^, 1970), mitochondria, chloroplasts, and possibly other organelles of eukaryotic cells descended from formerly free-living prokaryotes (Woese^16^, 1977; Lopez-Garcia et al.^17^, 2017; López-García and Moreira^18^, 2020; Zachar and Boza^19^, 2020; Zachar and Szathmáry^20^, 2017; Zachar^22^, 2022). During integration of mitochondria into an ancestor of eukaryotic cell, a new organism and a new level of complexity (multi-level selection) emerges from free-living and independently reproducing cells, and hence is one of the best examples of a major evolutionary transition (Szathmary^21^, 2015; Smith and Szathmary^6^, 1997). The establishment of a permanent and obligate coexistence of individuals that were once capable of independent existence played a crucial role in eukaryogenesis and organellogenesis.

One commonly proposed ecological dependence between species is syntrophy, i.e., one species living off the metabolic products of another species (Lopez-Garcia et al.^17^, 2017; López-García and Moreira^18^, 2020; Zachar and Boza^19^, 2020; Zachar and Szathmáry^20^, 2017). In this paper, we consider the resident ecological system of two species in which the host ancestor species produces the “food” of the symbiont ancestor species while the host itself is neither benefited nor harmed. Thereby the association can be called “syntrophic commensalism”. It is highly unlikely that the ancestral host and symbiont, at their very first interaction and onset of their association, turned out to be immediately mutualistic with compatible genome setups. Hence, we argue that it is plausible to consider the association to be a commensalistic one at the beginning with the potential to eventually evolve to a parasitic or even mutualistic one. Nevertheless, reciprocative metabolic exchange due to complementation (mutual or cyclic syntrophy) is assumed to be a key factor in the establishment of microbial associations and formation of microbial consortia, biofilms, and mats. However, in our model, we consider linear syntrophy rather than complex mutual syntrophy. It is widely assumed that interactions based on syntrophy might have played a significant part in the emergence of major endosymbiotic transitions and origin of mitochondria (Zachar and Szathmary^20^, 2017; Zachar et al.^13^, 2018; Lopez-Garcia et al.^17^, 2017; López-García and Moreira^18^, 2020; Zachar and Boza^19^, 2020). Various hypotheses of eukaryogenesis assume syntrophic interactions (Zachar and Szathmary^20^, 2017) (Hydrogen hypothesis, Sulphur hypothesis, etc.) between once independent organisms. The main difference between the phagocytosis hypothesis and syntrophy hypothesis is that by phagocytosis endosymbiosis is evolved directly while by syntrophy ecto-symbiosis is evolved first. However, in both scenarios the long-term association of the two different species evolved with vertical transmission.

Acknowledging the fact that the initial interactions and conditions of cooperative behaviour are widely debated, we narrow our scope to a single mechanism that could set off such an association. We hypothesize that the very first step in the evolution of cooperative behaviour between free-living organisms could be a single mutation in the symbiont genome that resulted in the formation of a new protein that is capable of latching onto the external surface of the host (pre-eukaryotes). We believe that it is highly likely that this single mutation could be the first step in the evolution of symbiosis rather than complex combinations of several mutations that could potentially lead to phagocytosis or any other mechanisms. However microbial syntrophy cannot explain how one partner gets inside the other (Zachar and Szathmary^20^, 2017) and we do suggest that development of phagocytotic features could be a later step in the process but not the first transition. In our model, we focus on this mutated symbiont ancestor that has the potential to form an ecto-symbiotic linkage with the host ancestor. The formation of this consortium is beneficial to the symbiont in the sense that it gains more and straightforward access to the substrate from the host. Clearly, in this scenario the ecto-symbiosis evolves, and the vertical transmission also immediately follows (assuming during the proliferation of the host ancestor the mutant symbiont is not lost). Here we introduce an ecological coevolutionary model, and we are interested in whether this mutant can invade into the resident system.

The successful invasion of a stable N-species resident system by mutant phenotypes has been discussed in (Cressman and Garay^23^, 2003; Garay and Varga^24^; 2000; Hofbauer and Sigmund^25^, 1998). In this paper, we are interested in the outcome of evolution by which the resident phenotype of one of the species (symbiont) dies out and is replaced by the mutant phenotype of the same species. We consider that mutation is rare enough and that multiple mutant phenotypes either in the same species or in several species does not occur at the same time. Evolutionary substitution occurs if the mutant phenotype can invade a stable equilibrium of the monomorphic N-species resident system (with all species) and the system evolves to a stable equilibrium of the N-species resident-mutant coevolutionary system (also with all species present) but the one species has only the mutant phenotype (Cressman et al.^26^, 2020; Cressman and Garay^27^, 2003).

In other words, the requirements for evolutionary substitution are:

1. The resident system with no mutant phenotype is stable, i.e, there is species coexistence in the resident system. Mathematically, there exists a locally asymptotically stable positive equilibrium to the resident system.
2. The mutant phenotype (of exactly one species) can invade, i.e, the resident-only equilibrium of the resident-mutant system is not a locally asymptotically stable equilibrium.
3. The resident-mutant system must evolve to a locally asymptotically stable equilibrium with all species present, but species which underwent the mutation has only mutant phenotype.

We investigate the ecological and evolutionary stability of inter-species microbial interactions in the framework of an ecto-commensalistic association based on syntrophy using the above-mentioned concept. The aim of this paper is to obtain a set of sufficient conditions which ensures the existence of the positive equilibrium point of the coevolutionary resident-mutant system with the mutant phenotype, which is locally stable. We thereby analyze the sufficient conditions needed for pairwise interactions to emerge and stabilize in a system. To the best of our knowledge, this is the first time that a commensalistic model based on syntrophy hypothesis is considered in the framework of coevolutionary dynamics.

## Model

### Resident syntrophic ecological system

The first step in formulating a model for the commensalistic cross-feeding consists of setting up a system for the ecological dynamics of a monomorphic resident population consisting of a single phenotype of each species. Here we start from two free-living species that are in commensalistic association based on syntrophy. We assume species-X (host, pre-eukaryotes) is much bigger in size than species-Y (symbiont) and the metabolic product of X is consumed by Y.

The following basic assumptions are made to formulate the mathematical model corresponding to the resident system with free-living X and Y species:

a. Both the species are in commensal interaction, i.e., the free-living Y is a beneficiary of the free-living X, whereas X is neither benefited nor harmed by Y. Also, Y cannot acquire any sort of benefit in the form of resources for its survival without the presence of X (obligate relationship). Therefore, the model may be called as the “**syntrophic commensalism model**” based on nutrition.
b. The model consists of dynamics of resource concentration *w*(*t*) ∈ ℝ_≥0_ of the metabolic product of host species and dynamics of the two populations: free-living X and free-living Y whose densities are denoted by *x*(*t*) ∈ ℝ_≥0_ and *y*(*t*) ∈ ℝ_≥0_ at time t, respectively.
c. The concentration of substrate consumed by X is constant in unit time, i.e., X cannot decrease the concentration of available substrate on the nutrient surface from which X acquires its “food”.
d. There is no physical contact between X and the resident Y. Thus, the concentration of metabolic product w available to Y for consumption is small due to the dilution effect. The metabolic product w of X is diluted in the volume of the habitat or medium.
e. All the consumptions are linear (see Materials and Methods for more details).
f. There is no competition between X and Y, for instance, for space, etc.
g. In the presence of abundant resources, both the population densities *x*(*t*) and *y*(*t*) grow at the Malthusian growth rates α_*X*_ and α_*Y*_ respectively (see Materials and Methods for more details). One of the novelties of our work is the introduction of a new Malthusian parameter definition, according to which 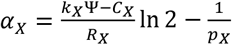 and 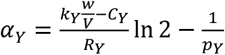.

Using the above basic assumptions, we propose the following deterministic model for the resident system using a set of non-linear (autonomous) differential equations:

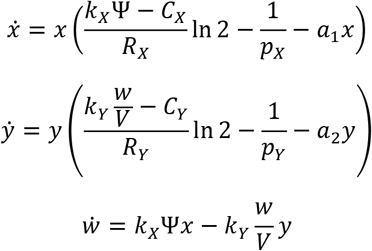

where all the parameters *k*_*X*_ ∈ (0, 1), Ψ ∈ ℝ_+_, *C*_*X*_ ∈ (0, 5), *R*_*X*_ ∈ (0, 5), *p*_*X*_ > 1, *a*_1_∈ ℝ_+_, *a*_2_ ∈ ℝ_+_, *k*_*Y*_ ∈ (0, 1), *C*_*Y*_ ∈ (0, 2), *R*_*Y*_ ∈ (0, 1), *p*_*Y*_ > 1and *V* ∈ ℝ_+_ are strictly positive real and are defined as in Table 1 (see materials and Methods for details).

**Table1:** Notations used in the main text and their meanings.

| Notations | Meaning |
| --- | --- |
| $x(t)$ | Density of free-living host |
| $y(t)$ | Density of free-living resident symbiont |
| $w(t)$ | Concentration of metabolic product of host species in the habitat |
| $z(t)$ | Density of consortium |
| $u(t)$ | Density of free-living mutant symbiont |
| $\Psi$ | Concentration of nutrient source available to host |
| $V$ | Volume of habitat or medium |
| $a_1$ | Competition within host species |
| $a_2$ | Competition within symbiont species |
| $k_X$ | Rate of consumption of $\Psi$ by host species |
| $k_Y$ | Rate of consumption of w by symbiont species |
| $C_X$ | Cost of living of host species |
| $C_Y$ | Cost of living of resident symbiont species |
| $R_X$ | Cost of reproduction of host species |
| $R_Y$ | Cost of reproduction of resident symbiont |
| $p_X$ | Death rate of host species |
| $p_Y$ | Death rate of symbiont species |
| $\alpha_X$ | Malthusian growth rate of host species |
| $\alpha_Y$ | Malthusian growth rate of resident symbiont |
| $C_U$ | Cost of living of mutant symbiont species |
| $R_U$ | Cost of reproduction of mutant symbiont |
| $\beta$ | Rate of interaction of host species and mutant symbionts |
| $h$ | Average number of mutant symbionts on the surface of a host in consortium |
| $\chi$ | Rate of inclusion of mutant symbionts into the habitat (in-consortium to free-living state) |
| $\alpha_U$ | Malthusian growth rate of mutant symbiont species |

### Coevolutionary system of ecto-symbiont

Now we consider a system (as shown in Figure1) which, in addition to the resident system, contains symbionts with a spatial mutation. The mutant symbiont U (with density u(t) ∈ ℝ_≥0_ when in free-living state) has a protein to join to the surface of its “host” (species X). For this we assume that there is a membrane protein (or membrane “carbon hydrogen”) of X, to which U can connect using its protein. We presume that this single mutation is the first step in the evolution of ecto-symbiosis. Thereby we now have a population comprising of host X in ecto-symbiotic association with U. Let us denote the ecto-symbiont “consortium or alliance (Z)” density by *z* ∈ ℝ_≥0_. It is important to note that the single mutation results in two extra dimensions in the coevolutionary dynamics of the entire system.

**Figure 1:**
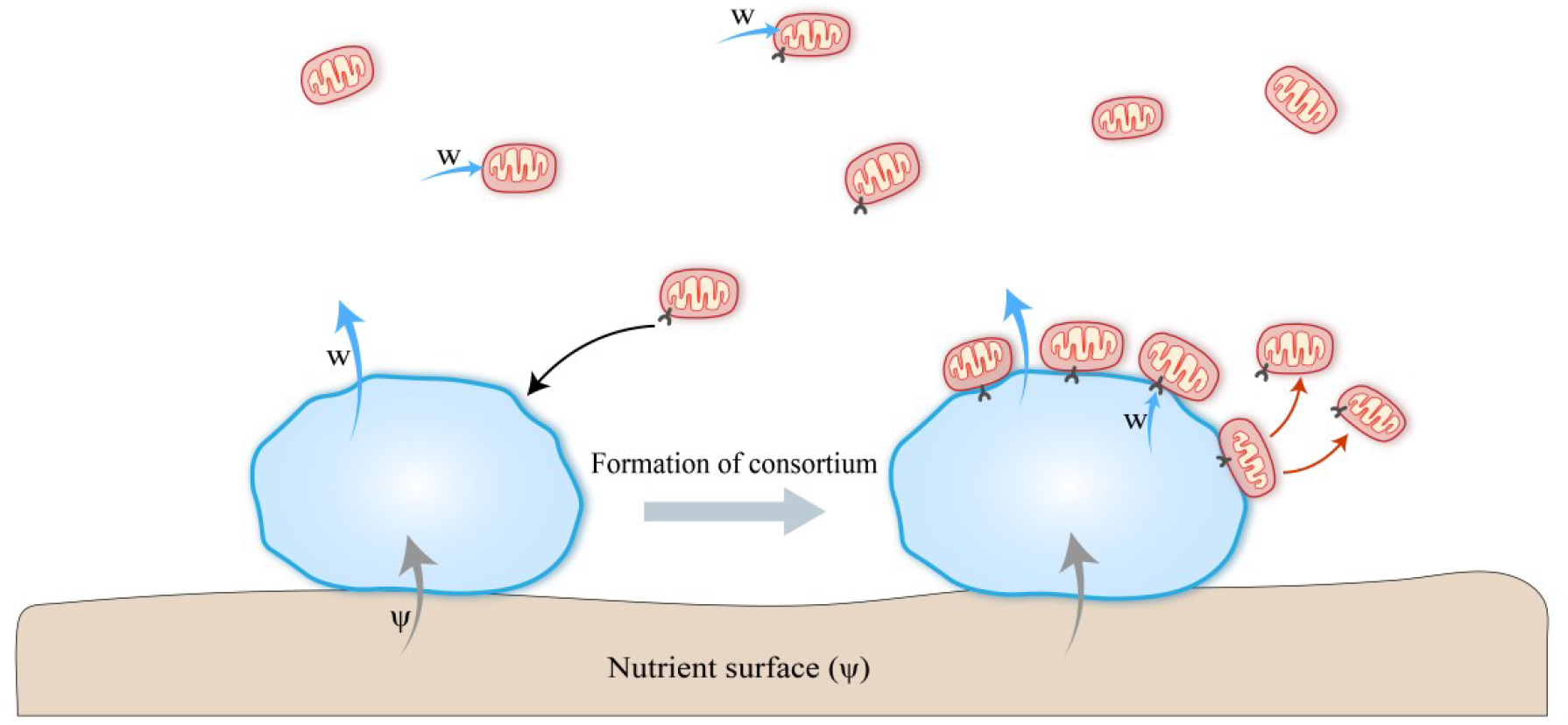
Schematic diagram of the resident-mutant coevolutionary system with host (blue) and symbiont (red) species. Grey arrows represent flow of Ψ, blue arrows represent flow of w (metabolic product of host species), black arrow represents formation of physiological connection between mutant symbiont and host’s external surface, and red arrows represent reproduction of mutant symbionts on host surface.

The following basic assumptions are made to formulate the mathematical model corresponding to the coevolutionary system with both resident and mutant phenotypes of the symbiont:

1. The “consortium, Z” can proliferate in the sense that the bigger member of Z, i.e, species-X multiplies, and the symbiont (species-U) is vertically transmitted during reproduction of the “consortium”. In other words, during the proliferation of X in alliance, the symbiont in the association is not lost and both parent and offspring cells of X have ecto-symbionts on their surface.
2. One free living host can collect more than one mutant symbiont U. Let us denote the average number of individuals of U on one host Z as *h*.
3. There is no competition between X and Y or X and U. But there is competition between X and Z. Also, there is competition between free-living Y and free-living U for dissolved w.
4. We assume that the symbiont U does not alter any characteristics of the host.
  a. the consumption rate of substrate Ψ by X remains constant, i.e, *k*_*X*_ = *k*_*Z*_,
  b. the death rate of X remains constant i.e, *p*_*X*_ = *p*_*Z*_,
  c. cost of living of X remains constant, i.e, *C*_*X*_ = *C*_*Z*_,
  d. cost of reproduction of X remains constant, i.e, *R*_*X*_ = *R*_*Z*_,
  e. Observe that assumptions (a)-(d) imply that *α_X_* = *α_Z_*. All these suggest that our model is an example of commensalism wherein the host in the alliance gets neither benefit nor loss.
5. The free-living U consumes the dissolved w much alike free-living Y. The consumption rate of dissolved w by U is the same as that of Y, i.e, *k*_*Y*_ = *k*_*U*_. In other words, we assume that the connecting protein and the connection does not alter the consumption rate.
6. Once connected with a host, U can consume the metabolic product w directly from its host at the rate *k*_*U*_. *hk*_*U*_*k*_*X*_Ψ of the metabolic produce of a single Z is directly consumed by the U in alliance, i.e, there is no element of dilution and hence access to resource is better. Thus, only (1− *k*_*U*_*h*)*k*_*X*_Ψ*z* part of the metabolic product w of all Z is dissolved. If the U-in-alliance can consume all metabolic products w produced by an individual Z in each “alliance” then no amount of w is dissolved into the habitat. Once connected to the host surface, the direct consumption of w by U-in-alliance is so effective and enough that the consumption of dissolved w by U-in-alliance is neglected.
7. Since Y and U differ in only one protein and the production of each protein has extra cost, *C*_*Y*_ < *C*_*U*_, where *C*_*U*_ and *C*_*Y*_ are the cost of living of U and Y respectively.
8. Since the connected protein also needs more biomass to reproduce, we can say that *R*_*U*_ > *R*_*Y*_, where *R*_*U*_ and *R*_*Y*_ are the cost of reproduction of U and Y respectively.
9. For simplicity we assume competition independent death rates for free-living Y and free-living U, i.e, *p*_*Y*_ = *p*_*U*_. Also, there is no toxic effect due to abundance of any of the substrates.
10. Since free-living U only consume dissolved metabolic product w, its Malthusian growth rate is given by,

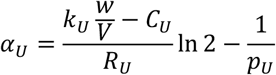
11. The encounter of free-living U and X is random, i.e., proportional to *xu*. New alliance is integrated by random encounters with probability *β* ∈ (0,1).
12. The addition of free-living U into the medium/ habitat can take place due to three phenomena.
  a. Z acts as an emitter of U: If U’s reproduction rate is high enough on the surface of the host, the Z individual produce free-living U at a constant rate *χ*_1_. When there is no free room (free surface) on the surface of the host, then there is an overflow of free-living U as the surface of the host is finite and thereby can get saturated. We assume U can reproduce faster than the host.
  b. Connection of U is not completely effective, i.e., the protein can detach from the surface of host. We assume there is only detachment of some U from the surface of host, but not all (This does not cause a reduction in density of Z). The detached U is added to free-living U population in the medium/ habitat at a constant rate *χ*_2_. Detachment rate is fixed, and this emission happens on a smaller time scale than the reproduction of the consortium.
  c. When a host-in-alliance dies, free-living U is added back to the medium or habitat at a constant rate 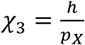. U-in-alliance goes back to its free-living state. Combining all additions of free-living U-species, we get, total addition = (*χ*_1_+ *χ*_2_ + *χ*_3_)*z*

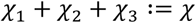

We assume total inclusion of free-living U into the medium is at a constant rate *χ* ∈ ℝ_+_. Note that high values of *χ* is indicative of high possibility of horizontal transmission of mutant symbionts as it increases the density of free-living U linearly.

From the above basic assumptions, we propose the following deterministic model for the dynamics of the resident-mutant coevolutionary system by using a set of non-linear (autonomous) differential equations:

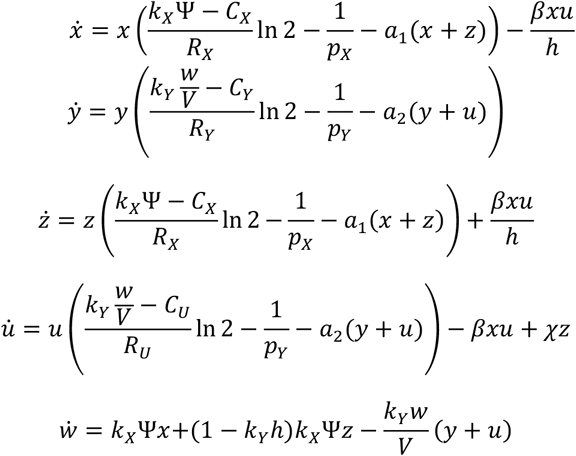

where all the parameters *k*_*X*_ ∈ (0, 1), Ψ ∈ ℝ_+_, *C*_*X*_ ∈ (0, 5), *R*_*X*_ ∈ (0, 5), *p*_*X*_ > 1, *a*_1_∈ ℝ_+_, *a*_2_ ∈ ℝ_+_, *k*_*Y*_ ∈ (0, 1), *C*_*Y*_ ∈ (0, 2), *R*_*Y*_ ∈ (0, 1), *C*_*U*_ ∈ (0, 2), *R*_*U*_ ∈ (0,1), *p*_*Y*_ > 1, *β* ∈ (0, 1), *h* ∈ ℝ_+_, *χ* ∈ ℝ_+_ and *V* ∈ ℝ_+_ are strictly positive real and are defined as in Table 1 (see Materials and Methods for more details).

## Results

The monomorphic resident system is analysed for the stability conditions, which is one of the primary objectives of our study. The system has two biologically feasible equilibrium points namely: the trivial equilibrium point *E*_0_(0, 0, 0) and the interior equilibrium point *E*_*R*_(*x*^*^, *y*^*^, *w*^*^). Both the equilibria exist unconditionally and *E*_0_(0, 0, 0) is unstable always and *E*_*R*_(*x*^*^, *y*^*^, *w*^*^) is locally asymptotically stable always (see SI (1) for proof). Thus, we have a special commensalistic dynamics in which if all the parameters are positive then we have locally asymptotically stable interior rest point in three-dimension. In other words, the resident system with no mutant phenotype is stable and there is coexistence of host and symbiont species in the absence of mutant phenotypes. Also, we can observe that the mutant-only equilibrium (*E*_*M*_(*z*^+^, *u*^+^, *w*^+^)) with resident host (in consortium) and mutant phenotype exists always and is locally asymptotically stable always (see SI (2) for proof).

**Figure 2(a):**
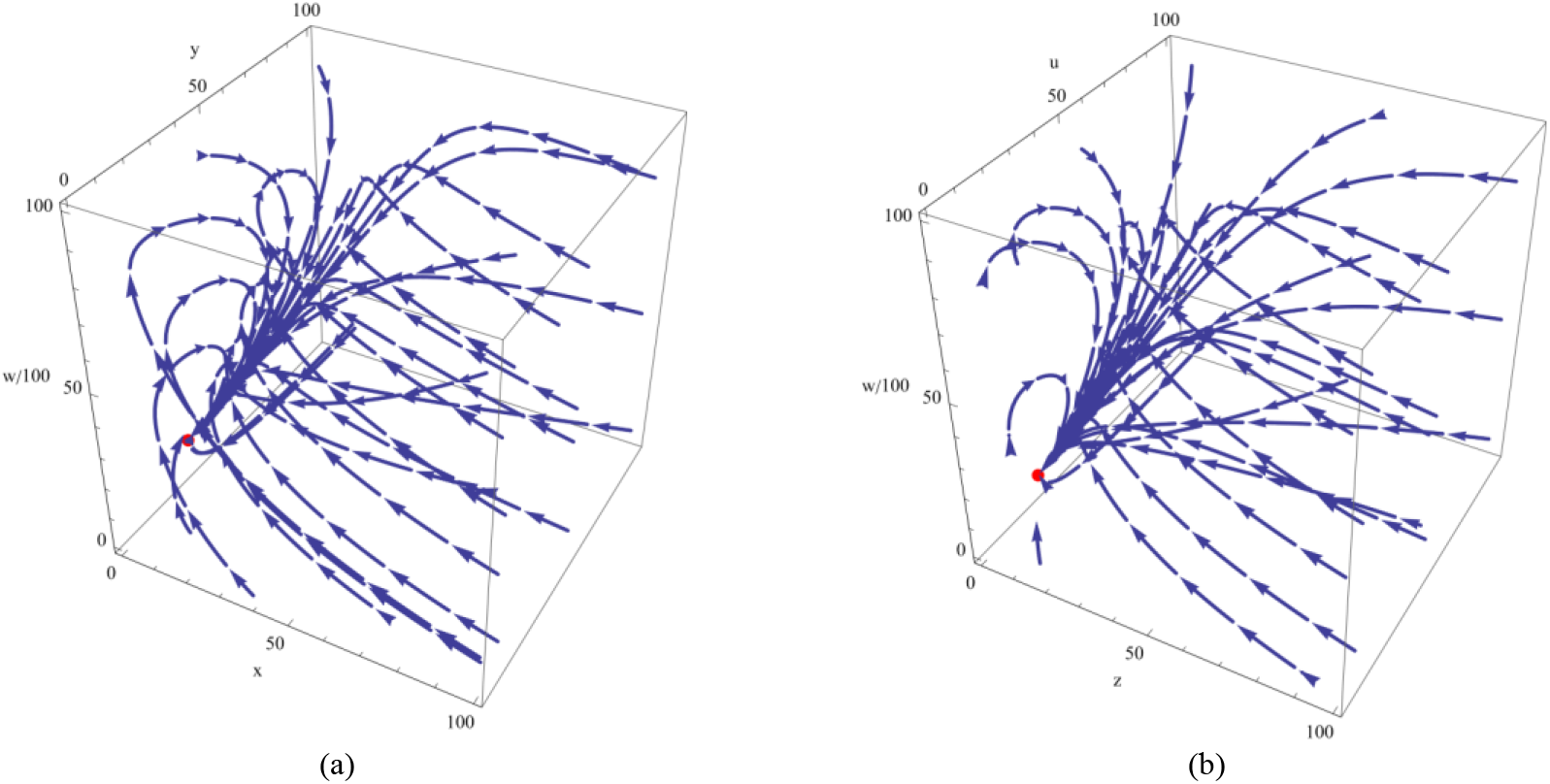
Phase-portrait of the resident-only system (3D) with trajectories tending to E_R_(x^*^, y^*^, w^*^) = (5.8, 22.4, 2076) when Ψ = 400, a_1_= 3, a_2_ = 2, k_X_ = 0.2, C_X_ = 4, R_X_ = 3, P_X_ = 10, k_Y_ = 0.1, R_Y_ = 0.3, C_Y_ = 1.2, P_Y_ = 3 and V = 10. Figure2(b): Phase-portrait of the mutant-only system (3D) with trajectories tending to E_M_(z^+^, u^+^, w^+^) = (5.8, 17.5, 1595) when Ψ = 400, a_1_= 3, a_2_ = 2, k_X_ = 0.2, C_X_ = 4, R_X_ = 3, P_X_ = 10, k_U_ = k_Y_ = 0.1, P_U_ = P_Y_ = 3, h = 4, χ = 30, R_U_ = 0.4, C_U_ = 1.3, and V = 10.

The five-dimensional coevolutionary resident-mutant system is analysed to find the sufficient conditions that would enable the ecological and evolutionary stability of the intimate consortium of two species that were once capable of independent mobility, which is the significant purpose of this paper. The resident-mutant system has four biologically feasible equilibria namely: *E*_0_(0,0,0,0,0), *E*_1_(*x*^*^, *y*^*^, 0, 0, *w*^*^), *E*_2_(0, 0, *z*^+^, *u*^+^, *w*^+^) and 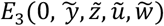. It is worthwhile to note that there does not exist an interior equilibrium point (*x, y, z, u, w*) of the five-dimensional coevolutionary dynamics such that x, y, z, u, w ∈ ℝ_+_. In other words, positive valued interior equilibria do not exist for the above-mentioned resident-mutant system and there are only boundary equilibria to the system.

The trivial equilibrium point *E*_0_(0,0,0,0,0) and resident trivial equilibrium point *E*_1_(x^*^, y^*^, 0, 0, w^*^) exist always unconditionally and are always unstable (see Theorem 2.1 in SI (2) for proof). Biologically, this means that the mutant phenotype of the symbiont can invade the resident system. Additionally, for the evolutionary substitution of the resident phenotype by the mutant phenotype of the symbiont, we need to check whether the resident-mutant system evolves to a locally asymptotically stable equilibrium with the mutant phenotype of the symbiont.

Mathematically, the mutant equilibrium point of the resident-mutant system *E*_2_(0,0, *z*^+^, *u*^+^, *w*^+^) exists unconditionally and is locally asymptotically stable if (Sufficient conditions that guarantees the invasion and subsequent fixation of the mutant system with the intimate consortium):

1. 1) 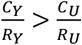 and
2. 2) 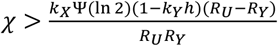

(See Theorem 2.2 in SI (2) for proof).

**Figure 3:**
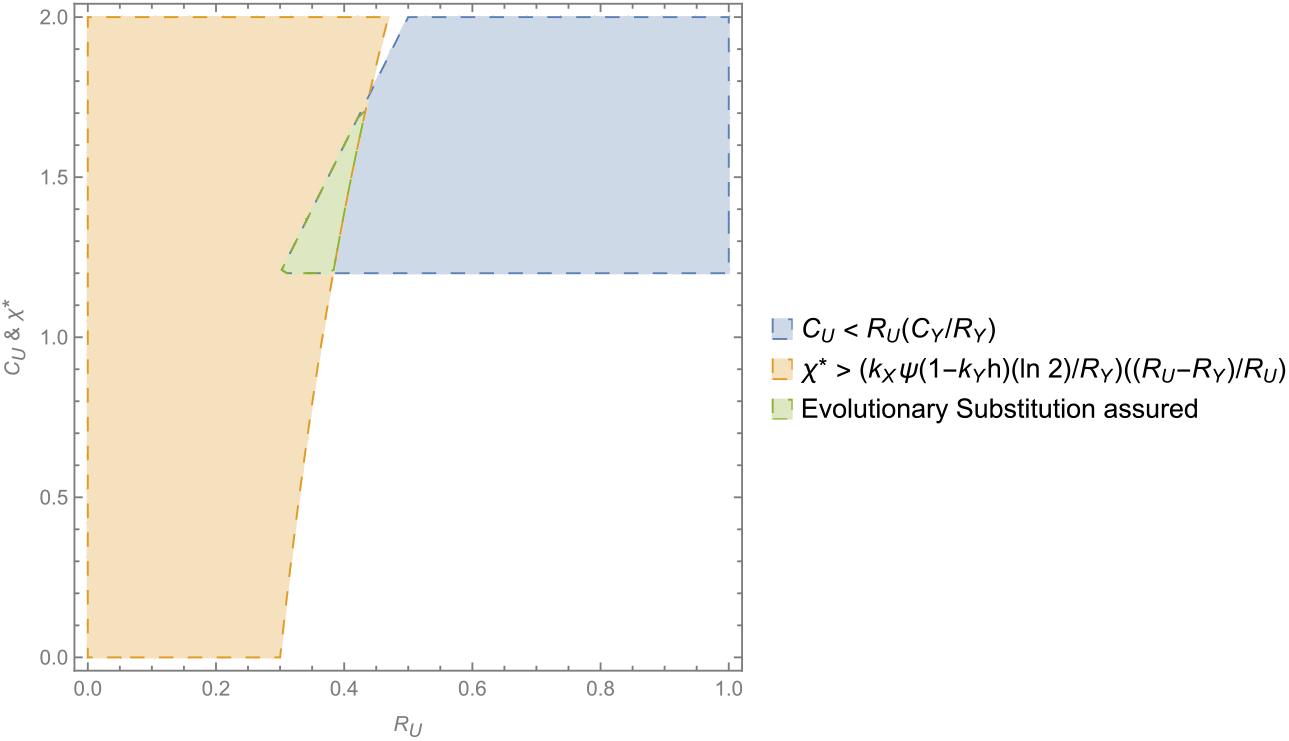
Region-plot with R_U_ on X-axis and C_U_ & scaled χ on Y-axis showing the sufficient conditions for evolutionary substitution of resident phenotype by mutant phenotype of the symbiont species. The values of all the other parameters are fixed and as in Figure2. Analyzing the plot, the minimum probability of evolutionary substitution can be obtained to be 0.011 (see SI (4) for details).

If the above conditions are satisfied, we observe evolutionary substitution, i.e, a mutant appears in the symbiont species, the resident phenotype in this species dies out, and the mutant coexists with the original phenotype of the host by forming the consortium. If these sufficient conditions hold, this rules out the existence of the equilibrium point 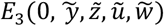. But if these conditions are not satisfied, then *E*_3_ can also exist and can also be locally asymptotically stable under some conditions. In other words, the mutant symbiont can also coexist with the resident symbiont along with the resident host. Conclusively, after successful invasion, the possible outcomes of evolution are evolutionary substitution and coexistence of resident and mutant phenotypes (polymorphism). However, we limit our focus only to the case of evolutionary substitution. From a biological perspective, the results of analysis suggest that the evolutionary substitution in our model happens when the cost of living to reproduction of the mutant phenotype is less than that of the resident phenotype of the symbiont and there is high possibility of horizontal transmission. This substitution of the resident phenotype by mutant phenotype would eventually mean the emergence and evolutionary stability of the consortium in the ecological system.

**Figure 4(a)-(b):**
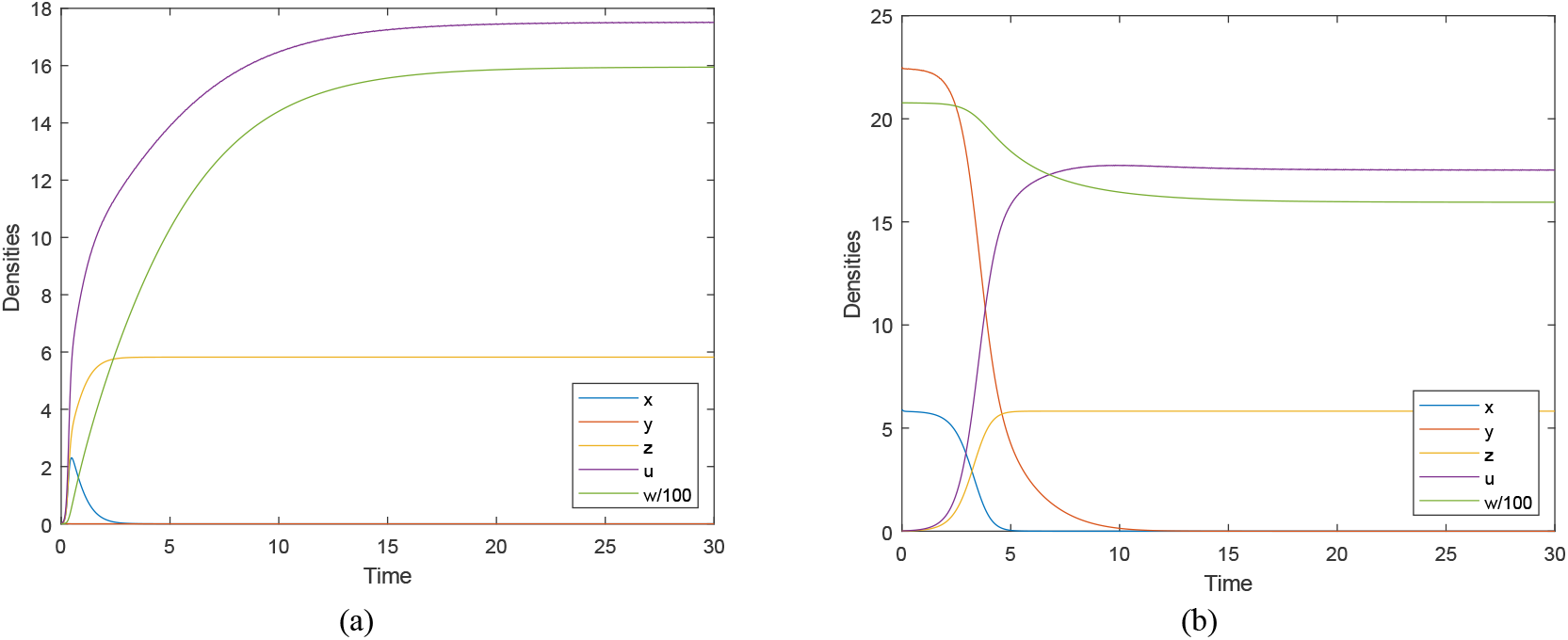
Time series plot of the resident-mutant system (5D) showing the stability of E_2_(0, 0, z^+^, u^+^, w^+^) = (0, 0, 5.8, 17.5, 1595) when Ψ = 400, a_1_= 3, a_2_ = 2, k_X_ = 0.2, C_X_ = 4, R_X_ = 3, P_X_ = 10, k_Y_ = 0.1, R_Y_ = 0.3, C_Y_ = 1.2, P_Y_ = 3, β = 0.8, h = 4, V = 10, C_U_ = 1.3, χ = **30, R**_**U**_ = **0. 4**; Figure4(a): For initial conditions x (0) = 0.01, y (0) = 0.01, z(0) = 0.01, u(0) = 0.01, w (0) = 0.01, we can observe that the dynamics attain stability at (0, 0, 5.8, 17.5, 1595). This validates that E_0_(0,0,0,0,0) is unstable and E_2_(0,0, z^+^, u^+^, w^+^) is stable. Figure4(b): For initial conditions x (0) = 5.9, y (0) = 22.5, z(0) = 0.01, u(0) = 0.01, w (0) = 2077, we can observe that the dynamics attain stability at (0, 0, 5.8, 17.5, 1595). This validates that E_1_(x^*^, y^*^, 0, 0, w^*^) = (5.8, 22.4, 0, 0, 2076) is unstable and E_2_(0,0, z^+^, u^+^, w^+^) is stable.

**Figure4(c)-(d):**
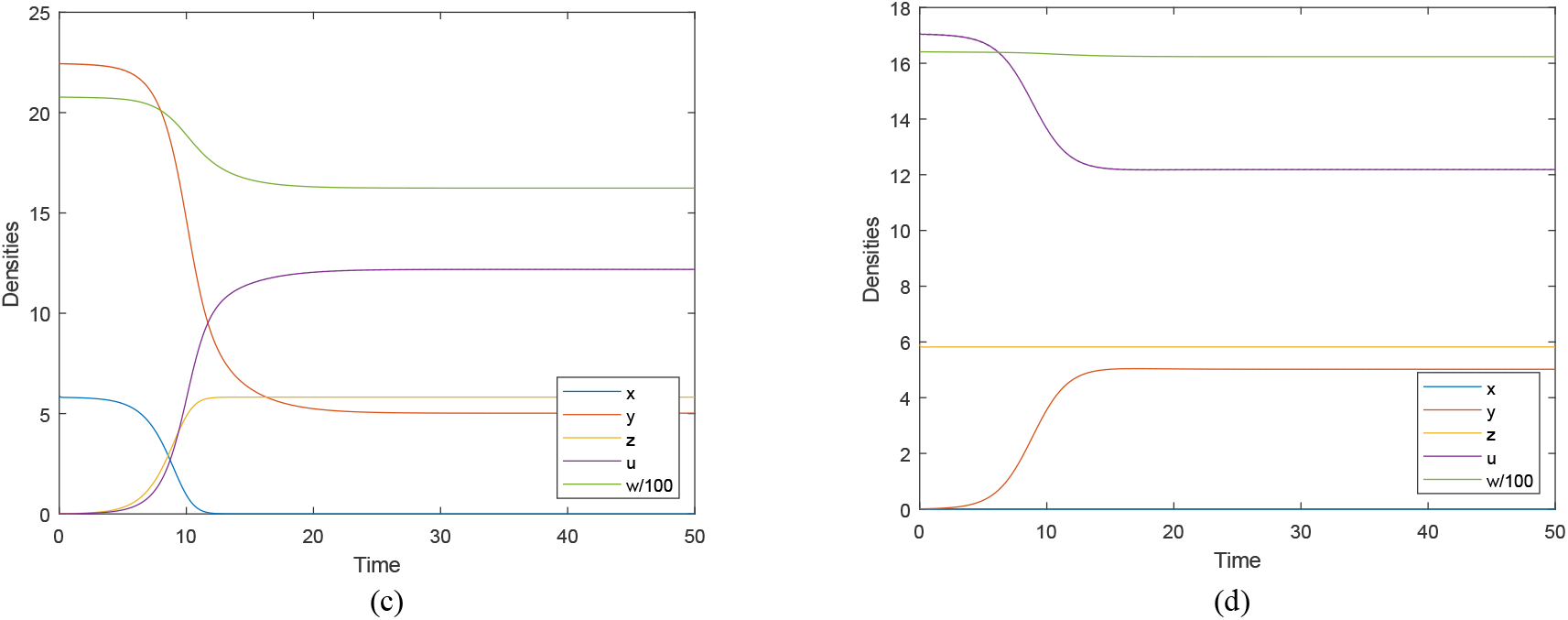
Time series plot of the resident-mutant system (5D) showing the stability of 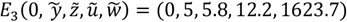,when Ψ = 400, a_1_= 3, a_2_ = 2, k_X_ = 0.2, C_X_ = 4, R_X_ = 3, P_X_ = 10, k_y_ = 0.1, R_Y_ = 0.3, C_Y_ = 1.2, P_Y_ = 3, β = 0.8, h = 4, V = 10, C_U_ = 1.3, χ = **5, R**_**U**_ = **0. 32**; Figure4(c): For initial conditions x (0) = 5.9, y (0) = 22.5, z(0) = 0.01, u(0) = 0.01, w (0) = 2077 we can observe that the dynamics attain stability at (0, 5, 5.8, 12.2, 1623.7). This validates that E_1_(x^*^, y^*^, 0, 0, w^*^) is unstable and 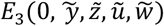 is stable; Figure4(d): For initial conditions x (0) = 0.01, y (0) = 0.01, z(0) = 5.9, u(0) = 17.1, w (0) = 1641, we can observe that the dynamics attain stability at (0, 5, 5.8, 12.2, 1623.7). This validates that E_2_(0, 0, z^+^, u^+^, w^+^) = (0, 0, 5.8, 17, 1640) is unstable and 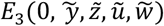 is stable. Note here that the values of z^+^, u^+^and w^+^ changes depending on change in the values of R_U_ and χ.

## Discussion

From an evolutionary perspective, beneficial (adaptive) mutations are very rare and hence it is reasonable to argue that the evolution is a step-by-step process. We started from a commensal ecological connection where the ancestral host’s metabolic product is food to the symbiont ancestor. We assume that only one mutation occurs, i.e, the symbionts are joined by a mutant membrane protein to the external surface of the host ancestor. Here the main point is that this mutation forms a new level of selection based on the fact that the symbiont can vertically transmit. Using theoretical models, we find that if the cost of living to reproduction of the mutant phenotype is less than that of the resident phenotype of the symbiont and there is high possibility of horizontal transmission, then the host-symbiont association or consortium is fixed by natural selection. Note that the additional costs required for living and reproduction of the mutant phenotype play a significant factor in fixation of consortium in the ecosystem. Even though both the respective costs of mutant phenotype are greater than that of the resident phenotype, it is worthwhile to note that it is the ratio of the costs that matters. After the fixation of host-symbiont association in the ecosystem, there are two different and broad directions for evolutionary mechanisms to proceed.

The first is that the association can evolve to be a mutualistic one in the sense that, at some point of time excess concentrations of the metabolic product of the host in its proximity turns out to be toxic to itself and the symbiont plays a significant role in reducing this concentration below a critical level and thereby helping the host in its survival. Another possibility is the evolution of cyclic-syntrophy, i.e., as a result of a mutation on the host, it can use at least one of the metabolic products of its ecto-symbiont, while the symbiont has not changed itself by other mutations. This is similar to the concept of evolutionary replacement discussed in Cressman and Garay^26^, 2020. Unlike evolutionary substitution, evolutionary replacement requires that there are mutant phenotypes in each of the species. Observe that in this scenario a costless mutualism can emerge because each member consumes the metabolic product of its partner. Moreover, since the host ancestor and symbiont ancestor are different species, we can assume that the metabolic ability of the species are quite different, and thereby the symbiont can potentially produce metabolic products that the host is incapable of producing itself and can use as food. In the case of mutualistic interaction, the vertical transmission is more evolutionarily rational (Bright and Bulgharesi^8^, 2010) since multilevel selection can enforce better association (i.e., associations have beneficial connection for both the host and its vertically transmitted symbiont, thus the reproduction rate of this association should be higher than other possibilities). Clearly, vertical transmission is more likely for the endo-symbiont than for the ecto-symbiont. Thus, the mutations which force the vertical transmission should invade into the population. For instance, suppose the ecto-symbiont can move into host cell by engulfement via phagocytosis or via slowly increasing surface contact during ecto-symbiotic syntrophy, or by bacterial invasion (Zachar and Szathmary^20^, 2017). After this the “consensus” of the originally almost independent metabolisms of the host and its endo-symbionts can lead to the emergence of endosymbiosis like the mitochondrial ancestor (Lopez-Garcia et al.^17^, 2017; López-García and Moreira^18^, 2020; Zachar and Boza^19^, 2020).

The other broad evolutionary possibility or direction is that by a new mutation the ecto-symbionts can kill their host and consume the host biomass for its own good. In this way the pathogen evolution is started. We note that, in our model, the symbiont has been emitted by consortium, thus the horizontal transmission is also possible. In summary, we stick to the idea that one of the important steps in the evolution of the symbiosis could be a mutation which results in vertical transmission. The vertical transmission form associations and the evolutionary success of vertically transmitted symbiont strictly depends on that of the host in the association.

## Materials and Methods

### General model for the Malthusian growth rate

We introduce our formulation of the growth rate of the species using a generalized model of ageless monas (unicellular organism) based on its life-history. We assume that the monas X cannot phagocytose and that it does not age. Species X consumes substrate Ψ as “food” and for simplicity we assume that the concentration of Ψ does not change in unit time, i.e. Ψ consumed by X is immediately replaced. If the monas has collected enough biomass, i.e., when it reaches a threshold or critical quantity (cost of reproduction), then it proliferates. For the simplicity we assume that during proliferation the organism does not die, and that the time duration of proliferation can be neglected. In other words, time duration of collection of biomasses for reproduction is longer than the time duration of the proliferation.

Assume that the consumption is linear, deterministic, continuous, and uniform in time with rate *l*, and the death rate of the monas is *p*. Now we are looking for the Malthusian growth rate of this ageless lineage. If the monas lives till time *R*/*l*, it proliferates. This happens with probability π ≔ *e*^−*R*/*pl*^. Thus, the distribution of its lifespan is exponential with parameter *p* and truncated at *R*/*l*. The expectation of the offspring sizes up to time *t* is equal to 0, if *t* < *R*/*l* and 2π, if *t* ≥ *R*/*l*. This branching process is supercritical if and only if 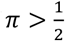, that is, 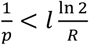.

The reproduction measure is degenerate: it puts weight 2π at point *R*/*l*. Therefore, the Malthusian equation is 2π*e*^−*αR*/*l*^ = 1, from which the Malthusian parameter is

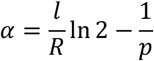

### Malthusian growth rate of host species

The Malthusian growth rate of the host species depends on the following factors:

#### 1. Consumption

For the simplicity we assume the “food or resource” concentration Ψ ∈ ℝ_+_ is constant in unit time. For instance, species X (host) lives on a nutrient surface Ψ that is continuously replenished. We assume the feeding is linear. Let *k*_*X*_ ∈ (0,1) be the consumption rate of Ψ by species X, then the consumption in unit time is,

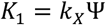

#### 2. Fertility based on biomass (energy) balance

Let us denote the cost of living of X by *C*_*X*_ ∈ (0, 5). Thus, the net biomass available for reproduction of X during unit time is,

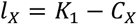

#### 3. Cost of reproduction

Denote *R*_*X*_ ∈ (0, 5) as the cost of reproduction, i.e., biomass *R*_*X*_ is the threshold needed for reproduction.

#### 4. Death rate

We assume that during unit time the death rate of each X individual is fixed i.e., independent of the concentration of metabolic product of X and is denoted by 1/*p*_*X*_, where *p*_*X*_ > 1and 1/*p*_*X*_ ∈ (0,1).

Substituting all the parameters into the equation of Malthusian parameter for species X,

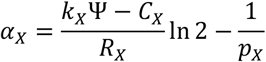

### Malthusian growth rate of symbiont species

The Malthusian growth rate of the symbiont species depends on the following factors:

#### 1. Consumption

Since species Y (symbiont) consumes the metabolic product of species X, the stoichiometry of the conversion Ψ → A + *w* is important. A is the metabolic product utilized by X about which we are not concerned. *w* is the metabolic product of X that gets dissolved in the surrounding habitat and is eventually utilized by species Y. For the simplicity we assume X makes exactly one molecule of *w* from one molecule of Ψ it consumes. In other words, concentration of *w* is proportional to the consumption of Ψ by X. We assume the consumption of Y is linear. Let *k*_*Y*_ ∈ (0,1) be the consumption rate of *w* by Y in unit time. Denote by *V* the volume of the habitat. Note that 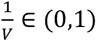 is representative of the dilution rate of *w*. As the volume of habitat increases, the concentration of dissolved *w* available to Y for consumption decreases. Thus, when the actual concentration is 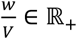, the “food” consumed by Y is given by,

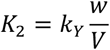

Since *w* dissolved in the habitat is in proportion to the consumption of food by X, the concentration of *w* in the habitat increases by *k*_*X*_Ψ*x* in unit time. Indeed, all X in the habitat collectively can consume *k*_*X*_Ψ*x* molecules of Ψ. Moreover, Y can consume *k* differential equation, 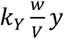, thus the concentration of w in the habitat is given by the following differential equation,

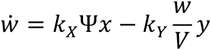

Observe here we assume the metabolic product of X (which is the food for Y) is chemically stable. In other words, the degradation time of *w* is much longer than the reproduction time of Y, thus Y can consume *w* before it is chemically degraded or transformed.

#### 2. Biomass (energy) balance

The cost of living for Y is denoted by *C*_*Y*_ ∈ (0, 2). Thus, the collected biomass for the reproduction of species Y in the unit time is,

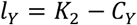

#### 3. Cost of reproduction

Denote *R*_*Y*_ ∈ (0, 1) the cost of reproduction of symbiont species, i.e., biomass *R*_*Y*_ is the threshold needed for reproduction.

#### 4. Death rate

We assume that during unit time, the death rate of each Y individual is fixed, denoted by *p*_*Y*_ > 1 and 1/*p*_*Y*_ ∈ (0,1), i.e., death rate does not depend on concentration of *w*.

Substituting all the parameters into the equation of Malthusian parameter for species Y, we get,

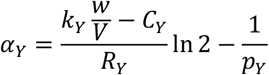

## Acknowledgements

This project has received funding from the European Union’s Horizon 2020 Research and Innovation Programme under the Marie Skłodowska-Curie grant agreement number 955708.

