## Supplementary Information for "Genesis of Ecto-symbiotic features based on Commensalistic Syntrophy"

### (SI 1) Equilibria and Local stability analysis of the resident system

Based on the description of our resident model, we have the following three-dimensional dynamical system:

$$\dot{x} = x\left(\frac{k_X\Psi - C_X}{R_X}\ln 2 - \frac{1}{p_X} - a_1 x\right)$$

$$\dot{y} = y\left(\frac{k_Y\frac{w}{V} - C_Y}{R_Y}\ln 2 - \frac{1}{p_Y} - a_2 y\right)$$

$$\dot{w} = \left(k_X\Psi x - k_Y\frac{w}{V}y\right)$$

where all the parameters $k_X \in (0,1)$, $\Psi \in \mathbb{R}_+$, $C_X \in (0,5)$, $R_X \in (0,5)$, $p_X > 1$, $a_1 \in \mathbb{R}_+$, $a_2 \in \mathbb{R}_+$, $k_Y \in (0,1)$, $C_Y \in (0,2)$, $R_Y \in (0,1)$, $p_Y > 1$ and $V \in \mathbb{R}_+$ are strictly positive real and are defined as in Table 1 of the main text.

Let us first redefine the system as following:

$$\dot{x} = x(\alpha - a_1 x)$$

$$\dot{y} = y(\eta w - \rho - a_2 y)$$

$$\dot{w} = \phi x - \xi w y$$

where all the parameters $\alpha, \eta, \rho, \phi, \xi$ are strictly positive and are defined as follows:

$$\alpha = \alpha_X = \frac{k_X\Psi - C_X}{R_X}\ln 2 - \frac{1}{p_X}$$

$$\eta = \frac{k_Y}{VR_Y}\ln 2, \qquad \rho = \frac{C_Y}{R_Y}\ln 2 + \frac{1}{p_Y}$$

$$\phi = k_X\Psi, \qquad \xi = \frac{k_Y}{V}$$

We consider ecologically meaningful initial conditions $x(0) > 0, y(0) > 0, w(0) > 0$.

Next, we investigate the dynamics of the system in $\mathbb{R}_+^3$. The system has two biologically feasible equilibria namely: $E_0(0,0,0)$ and $E_R(x^*, y^*, w^*)$ where,

$$x^* = \frac{\alpha}{a_1}$$

$$y^* = \frac{\sqrt{\rho^2 + \frac{4a_2\alpha\eta\phi}{a_1\xi}} - \rho}{2a_2}$$

$$w^* = \frac{\alpha\phi}{a_1\xi y^*} = \frac{\sqrt{\rho^2 + \frac{4a_2\alpha\eta\phi}{a_1\xi}} + \rho}{2\eta}$$

Clearly both the equilibria $E_0(0,0,0)$ and $E_R(x^*, y^*, w^*)$ exist unconditionally.

Now, we investigate the stability of the resident system at both the equilibria.

The Jacobian matrix $J(x, y, w)$ of the dynamical system at any arbitrary point $(x, y, w)$ is given by,

$$J_{(x,y,w)} = \begin{pmatrix} \alpha - 2a_1x & 0 & 0 \\ 0 & \eta w - \rho - 2a_2y & \eta y \\ \phi & -\xi w & -\xi y \end{pmatrix}$$

Clearly, $E_0(0,0,0)$ is unstable as one eigenvalue of $J_{(0,0,0)}$ is positive.

**Theorem 1.1: (Local stability of the resident-only positive equilibrium)**

The interior equilibrium $E_R(x^*, y^*, w^*)$ of the resident system is locally asymptotically stable unconditionally.

Proof:

Jacobian matrix at $(x^*, y^*, w^*)$:

$$J_{(x^*,y^*,w^*)} = \begin{pmatrix} \alpha - 2a_1x^* & 0 & 0 \\ 0 & \eta w^* - \rho - 2a_2y^* & \eta y^* \\ \phi & -\xi w^* & -\xi y^* \end{pmatrix}$$

According to Routh Hurwitz criterion, the eigenvalues of the Jacobian matrix are negative for the following conditions:

1) $\operatorname{tr} J_{(x^*,y^*,w^*)} < 0$ ,
2) $\det J_{(x^*,y^*,w^*)} < 0$,
3) $\Lambda = \det\begin{pmatrix} J_{11} & 0 \\ 0 & J_{22} \end{pmatrix} + \det\begin{pmatrix} J_{11} & 0 \\ J_{31} & J_{33} \end{pmatrix} + \det\begin{pmatrix} J_{22} & J_{23} \\ J_{32} & J_{33} \end{pmatrix} > 0$
4) $\operatorname{tr} J_{(x^*,y^*,w^*)}\Lambda < \det J_{(x^*,y^*,w^*)}$

$$J_{(x^*,y^*,w^*)} = \begin{pmatrix} \alpha - 2a_1x^* & 0 & 0 \\ 0 & \eta w^* - \rho - 2a_2y^* & \eta y^* \\ \phi & -\xi w^* & -\xi y^* \end{pmatrix} = \begin{pmatrix} J_{11} & 0 & 0 \\ 0 & J_{22} & J_{23} \\ J_{31} & J_{32} & J_{33} \end{pmatrix}$$

$$J_{11} = \alpha - 2a_1x^* = -\alpha < 0$$

$$J_{33} = -\xi y^* < 0$$

$$J_{22} = \eta w^* - \rho - 2a_2 y^* = \eta \frac{\sqrt{\rho^2 + \frac{4a_2\alpha\eta\phi}{a_1\xi}} + \rho}{2\eta} - \rho - 2a_2 \frac{\sqrt{\rho^2 + \frac{4a_2\alpha\eta\phi}{a_1\xi}} - \rho}{2a_2}$$

$$= \frac{\sqrt{\rho^2 + \frac{4a_2\alpha\eta\phi}{a_1\xi}} + \rho}{2} - \rho - \sqrt{\rho^2 + \frac{4a_2\alpha\eta\phi}{a_1\xi}} + \rho < 0$$

All diagonal elements of $J_{(x^*,y^*,w^*)}$ are negative and hence $\mathrm{tr}\, J_{(x^*,y^*,w^*)} < 0$.

Also, since $J_{23} = \eta y^* > 0$ and $J_{32} = -\xi w^* < 0$, we get $J_{22}J_{33} - J_{23}J_{32} > 0$.

Therefore, $\det J_{(x^*,y^*,w^*)} = J_{11}(J_{22}J_{33} - J_{23}J_{32}) < 0$ and

$$\Lambda = \det\begin{pmatrix} J_{11} & 0 \\ 0 & J_{22} \end{pmatrix} + \det\begin{pmatrix} J_{11} & 0 \\ J_{31} & J_{33} \end{pmatrix} + \det\begin{pmatrix} J_{22} & J_{23} \\ J_{32} & J_{33} \end{pmatrix} > 0.$$

Similarly, we can prove that,

$$\mathrm{tr}\, J_{(x^*,y^*,w^*)}\Lambda - \det J_{(x^*,y^*,w^*)} = (J_{11} + J_{22} + J_{33})(J_{11}J_{22} + J_{11}J_{33} + J_{22}J_{33} - J_{23}J_{32}) - J_{11}(J_{22}J_{33} - J_{23}J_{32}) < 0$$

as all the individual terms in the expansion are negative.

Alternative method:

The eigenvalues of $J_{(x^*,y^*,w^*)}$ are

$$J_{11}, \frac{1}{2}\left(J_{22} + J_{33} - \sqrt{J_{22}{}^2 + 4J_{23}J_{32} - 2J_{22}J_{33} + J_{33}{}^2}\right), \frac{1}{2}(J_{22} + J_{33} + \sqrt{J_{22}{}^2 + 4J_{23}J_{32} - 2J_{22}J_{33} + J_{33}{}^2})$$

All eigenvalues are negative as $J_{11}, J_{22}, J_{33} < 0$ and $J_{23}J_{32} < 0$.

We can observe that all the eigenvalues of the linearized system about $(x^*, y^*, w^*)$ are real and negative. Therefore, $(x^*, y^*, w^*)$ is locally asymptotically stable.

**(SI 2) Equilibria and Local stability analysis of the resident-mutant system**

Based on the description of our resident-mutant coevolutionary model, we have the following five-dimensional dynamical system:

$$\dot{x} = x\left(\frac{k_X\Psi - C_X}{R_X}\ln 2 - \frac{1}{p_X} - a_1(x+z)\right) - \frac{\beta x u}{h}$$

$$\dot{y} = y\left(\frac{k_Y\frac{w}{V} - C_Y}{R_Y}\ln 2 - \frac{1}{p_Y} - a_2(y+u)\right)$$

$$\dot{z} = z\left(\frac{k_X\Psi - C_X}{R_X}\ln 2 - \frac{1}{p_X} - a_1(x+z)\right) + \frac{\beta x u}{h}$$

$$\dot{u} = u\left(\frac{k_Y\frac{w}{V} - C_U}{R_U}\ln 2 - \frac{1}{p_Y} - a_2(y+u)\right) - \beta x u + \chi z$$

$$\dot{w} = k_X\Psi x + (1 - k_Y h)k_X\Psi z - k_Y\frac{w}{V}(y + u)$$

where all the parameters $k_X \in (0,1)$, $\Psi \in \mathbb{R}_+$, $C_X \in (0,5)$, $R_X \in (0,5)$, $p_X > 1$, $a_1 \in \mathbb{R}_+$, $a_2 \in \mathbb{R}_+$, $k_Y \in (0,1)$, $C_Y \in (0,2)$, $R_Y \in (0,1)$, $C_U \in (0,2)$, $R_U \in (0,1)$, $p_Y > 1$, $\beta \in (0,1)$, $h \in \mathbb{R}_+$, $\chi \in \mathbb{R}_+$ and $V \in \mathbb{R}_+$ are strictly positive real and are defined as in Table 1 of the main text.

Let us first redefine the system as following:

$$\dot{x} = x(\alpha - a_1 x - a_1 z - (\beta/h)u)$$

$$\dot{y} = y(\eta w - \rho - a_2 y - a_2 u)$$

$$\dot{z} = z(\alpha - a_1 x - a_1 z) + (\beta/h)xu$$

$$\dot{u} = u(\mu w - \sigma - a_2 y - a_2 u - \beta x) + \chi z$$

$$\dot{w} = \phi x + \kappa\phi z - \xi w y - \xi w u$$

where all the parameters $\alpha, \eta, \rho, \phi, \xi, \mu, \sigma, \kappa$ are strictly positive and are defined as follows:

$$\alpha = \frac{k_X\Psi - C_X}{R_X}\ln 2 - \frac{1}{p_X}$$

$$\eta = \frac{k_Y}{VR_Y}\ln 2, \qquad \rho = \frac{C_Y}{R_Y}\ln 2 + \frac{1}{p_Y}$$

$$\phi = k_X\Psi, \qquad \xi = \frac{k_Y}{V}$$

$$\mu = \frac{k_Y}{VR_U}\ln 2, \qquad \sigma = \frac{C_U}{R_U}\ln 2 + \frac{1}{p_Y}$$

$$\kappa = (1 - k_Y h)$$

Next, we investigate the stability conditions of the coevolutionary system in $\mathbb{R}^5_+$.

The system has four biologically feasible equilibria namely: $E_0(0,0,0,0,0)$, $E_1(x^*, y^*, 0, 0, w^*)$, $E_2(0,0,z^+,u^+,w^+)$ and $E_3(0, \tilde{y}, \tilde{z}, \tilde{u}, \tilde{w})$.

Note: We consider $x, y, z, u, w \in \mathbb{R}_+$ . Therefore, the only positive-valued equilibrium points of the 5-dimensional system are: $(x^*, y^*, 0, 0, w^*)$, $(0, 0, z^+, u^+, w^+)$ and $(0, \tilde{y}, \tilde{z}, \tilde{u}, \tilde{w})$. This fact rules out the possibility of existence of an interior rest point of the 5-dimensional coevolutionary dynamics $(x, y, z, u, w)$ such that $x, y, z, u, w \in \mathbb{R}_+$. In other words, positive valued interior equilibria do not exist for the above mentioned 5D system.

**Existence and Local Stability of $E_1(x^*, y^*, 0, 0, w^*)$**

Consider the equilibrium point of the redefined system $(x^*, y^*, 0, 0, w^*)$ where,

$$x^* = \frac{\alpha}{a_1}$$

$$y^* = \frac{\sqrt{\rho^2 + \frac{4a_2\alpha\eta\phi}{a_1\xi}} - \rho}{2a_2}$$

$$w^* = \frac{\alpha\phi}{a_1\xi y^*} = \frac{\sqrt{\rho^2 + \frac{4a_2\alpha\eta\phi}{a_1\xi}} + \rho}{2\eta}$$

As mentioned before, the resident equilibrium point of the system exists always unconditionally. We now conduct the stability analysis of this equilibrium point using Linearization.

**Theorem 2.1: (Local stability of $E_0(0,0,0,0,0)$ and $E_1(\mathrm{x}^*, \mathrm{y}^*, 0, 0, \mathrm{w}^*)$)**

$E_0(0,0,0,0,0)$ and $E_0(\mathrm{x}^*, \mathrm{y}^*, 0, 0, \mathrm{w}^*)$ are always unstable.

Proof:

The Jacobian matrix $J(x,y,z,u,w)$ of the dynamical system at any arbitrary point $(x,y,z,u,w)$ is given by,

$$J_{(x,y,z,u,w)} = \begin{pmatrix} \alpha - 2a_1x - a_1z - (\beta/h)u & 0 & -a_1x & -(\beta/h)x & 0 \\ 0 & \eta w - \rho - 2a_2y - a_2u & 0 & -a_2y & \eta y \\ -a_1z + (\beta/h)u & 0 & \alpha - a_1x - 2a_1z & (\beta/h)x & 0 \\ -\beta u & -a_2u & \chi & \mu w - \sigma - a_2y - 2a_2u - \beta x & \mu u \\ \phi & -\xi w & \phi\kappa & -\xi w & -\xi y - \xi u \end{pmatrix}$$

Clearly, $E_0(0,0,0,0,0)$ is unstable as one eigenvalue of $J_{(0,0,0,0,0)}$ is positive.

Jacobian matrix at $(x^*, y^*, 0, 0, w^*)$:

$$J_{(x^*,y^*,0,0,w^*)} = \begin{pmatrix} \alpha - 2a_1x^* & 0 & -a_1x^* & -(\beta/h)x^* & 0 \\ 0 & \eta w^* - \rho - 2a_2y^* & 0 & -a_2y^* & \eta y^* \\ 0 & 0 & \alpha - a_1x^* & (\beta/h)x^* & 0 \\ 0 & 0 & \chi & \mu w^* - \sigma - a_2y^* - \beta x^* & 0 \\ \phi & -\xi w^* & \phi\kappa & -\xi w^* & -\xi y^* \end{pmatrix}$$

For the local asymptotic stability of the equilibrium point, all eigenvalues of the Jacobian matrix must be negative.

Jacobian matrix at $(x^*, y^*, 0, 0, w^*)$:

$$J_{(x^*,y^*,0,0,w^*)} = \begin{pmatrix} \alpha - 2a_1x^* & 0 & -a_1x^* & -(\beta/h)x^* & 0 \\ 0 & \eta w^* - \rho - 2a_2y^* & 0 & -a_2y^* & \eta y^* \\ 0 & 0 & \alpha - a_1x^* & (\beta/h)x^* & 0 \\ 0 & 0 & \chi & \mu w^* - \sigma - a_2y^* - \beta x^* & 0 \\ \phi & -\xi w^* & \phi\kappa & -\xi w^* & -\xi y^* \end{pmatrix}$$

$$= \begin{pmatrix} J_{11} & 0 & J_{13} & J_{14} & 0 \\ 0 & J_{22} & 0 & J_{24} & J_{25} \\ 0 & 0 & 0 & J_{34} & 0 \\ 0 & 0 & J_{43} & J_{44} & 0 \\ J_{51} & J_{52} & J_{53} & J_{54} & J_{55} \end{pmatrix}$$

The eigenvalues are:

1) $J_{11}$,
2) $\frac{1}{2}\left(J_{44} - \sqrt{4J_{34}J_{43} + J_{44}^{\ 2}}\right)$
3) $\frac{1}{2}\left(J_{44} + \sqrt{4J_{34}J_{43} + J_{44}^{\ 2}}\right)$,
4) $\frac{1}{2}\left(J_{22} + J_{55} - \sqrt{J_{22}^{\ 2} + 4J_{25}J_{52} - 2J_{22}J_{55} + J_{55}^{\ 2}}\right)$,
5) $\frac{1}{2}(J_{22} + J_{55} + \sqrt{J_{22}^{\ 2} + 4J_{25}J_{52} - 2J_{22}J_{55} + J_{55}^{\ 2}})$

We can observe that the third eigenvalue is never negative since $J_{34}J_{43} > 0$.

Therefore $(x^*, y^*, 0, 0, w^*)$ is always unstable.

**Existence and Local Stability of $\boldsymbol{E_M(z^+, u^+, w^+)}$ and $\boldsymbol{E_2(0, 0, z^+, u^+, w^+)}$**

Let us now consider the equilibrium point of the redefined coevolutionary system $(0,0, z^+, u^+, w^+)$ where,

$$z^+ = \frac{\alpha}{a_1}$$

$$u^+ = \frac{\sqrt{\sigma^2 + \frac{4a_2\alpha(\mu\phi\kappa + \chi\xi)}{a_1\xi}} - \sigma}{2a_2}$$

$$w^+ = \frac{\alpha\phi\kappa}{a_1\xi u^+} = \frac{2\alpha\phi\kappa a_2}{a_1\xi\left(\sqrt{\sigma^2 + \frac{4a_2\alpha(\mu\phi\kappa + \chi\xi)}{a_1\xi}} - \sigma\right)}$$

$$= \frac{\phi\kappa\left(\sqrt{\sigma^2 + \frac{4a_2\alpha(\mu\phi\kappa + \chi\xi)}{a_1\xi}} + \sigma\right)}{2(\mu\phi\kappa + \chi\xi)}$$

The mutant rest point of the system exists always. We first conduct the stability analysis of the mutant-only equilibrium point in three-dimension.

Consider the following mutant-only system with only the mutant phenotype of the symbiont.

$$\dot{z} = z(\alpha - a_1 z)$$

$$\dot{u} = u(\mu w - \sigma - a_2 u) + \chi z$$

$$\dot{w} = \kappa\phi z - \xi w u$$

where all parameters are defined as in the five-dimensional coevolutionary system.

Jacobian matrix at $(z^+, u^+, w^+)$:

$$J_{(z^+,u^+,w^+)} = \begin{pmatrix} \alpha - 2a_1 z^+ & 0 & 0 \\ \chi & \mu w^* - \sigma - 2a_2 u^+ & \mu u^+ \\ \kappa\phi & -\xi w^+ & -\xi u^+ \end{pmatrix} = \begin{pmatrix} J_{11} & 0 & 0 \\ J_{21} & J_{22} & J_{23} \\ J_{31} & J_{32} & J_{33} \end{pmatrix}$$

$$J_{11} = \alpha - 2a_1 z^+ = -\alpha < 0$$

$$J_{33} = -\xi u^+ < 0$$

$$J_{22} = \mu w^+ - \sigma - 2a_2 u^+ = \mu \frac{\phi\kappa\left(\sqrt{\sigma^2 + \frac{4a_2\alpha(\mu\phi\kappa + \chi\xi)}{a_1\xi}} + \sigma\right)}{2(\mu\phi\kappa + \chi\xi)} - \sigma - 2a_2 \frac{\sqrt{\sigma^2 + \frac{4a_2\alpha(\mu\phi\kappa + \chi\xi)}{a_1\xi}} - \sigma}{2a_2}$$

$$= \frac{\mu\phi\kappa\left(\sqrt{\sigma^2 + \frac{4a_2\alpha(\mu\phi\kappa + \chi\xi)}{a_1\xi}} + \sigma\right) - 2(\mu\phi\kappa + \chi\xi)\sqrt{\sigma^2 + \frac{4a_2\alpha(\mu\phi\kappa + \chi\xi)}{a_1\xi}}}{2(\mu\phi\kappa + \chi\xi)} < 0$$

The eigenvalues of $J_{(z^+,u^+,w^+)}$ are

$$J_{11}\,, \frac{1}{2}\left(J_{22} + J_{33} - \sqrt{{J_{22}}^2 + 4J_{23}J_{32} - 2J_{22}J_{33} + {J_{33}}^2}\right), \frac{1}{2}(J_{22} + J_{33} + \sqrt{{J_{22}}^2 + 4J_{23}J_{32} - 2J_{22}J_{33} + {J_{33}}^2})$$

All eigenvalues are real and negative as $J_{11}, J_{22}, J_{33} < 0$ and $J_{23}J_{32} < 0$. Therefore, the interior equilibrium $E_M(z^+, u^+, w^+)$ of the mutant-only system is locally asymptotically stable always.

We now conduct the stability analysis of the mutant equilibrium point of the five-dimensional resident-mutant system.

**Theorem 2.2: (Local stability of the $E_2(0, 0, z^+, u^+, w^+)$)**

$E_2(0,0,z^+,u^+,w^+)$ is locally asymptotically stable if:

1) $\frac{C_Y}{R_Y} > \frac{C_U}{R_U}$ and
2) $\chi > \frac{k_X\Psi(\ln 2)(1-k_Y h)(R_U - R_Y)}{R_U R_Y}$

Proof:

Jacobian matrix at $(0,0,z^+,u^+,w^+)$:

$$J_{(0,0,z^+,u^+,w^+)} = \begin{pmatrix} \alpha - a_1 z^+ - (\beta/h)u^+ & 0 & 0 & 0 & 0 \\ 0 & \eta w^+ - \rho - a_2 u^+ & 0 & 0 & 0 \\ -a_1 z^+ + (\beta/h)u^+ & 0 & \alpha - 2a_1 z^+ & 0 & 0 \\ -\beta u^+ & -a_2 u^+ & \chi & \mu w^+ - \sigma - 2a_2 u^+ & \mu u^+ \\ \phi & -\xi w^+ & \phi\kappa & -\xi w^+ & -\xi u^+ \end{pmatrix}$$

The characteristic polynomial corresponding to the Jacobian matrix:

$$(\alpha - a_1 z^+ - (\beta/h)u^+ - \lambda)(\eta w^+ - \rho - a_2 u^+ - \lambda)(\alpha - 2a_1 z^+ - \lambda)[\lambda^2 + \lambda(2a_2 u^+ + \xi u^+ + \sigma - \mu w^+) + (\sigma\xi u^+ + 2a_2\xi {u^+}^2)] = 0$$

For the local asymptotic stability of the equilibrium point, all eigenvalues of the Jacobian matrix must be negative.

We can make the following observations:

1) $\alpha - a_1 z^+ - (\beta/h)u^+ < 0$

2) $\alpha - 2a_1 z^+ < 0$

3) $\sigma\xi u^+ + 2a_2\xi u^{+2} > 0$

4) $2a_2u^+ + \xi u^+ + \sigma - \mu w^+ > 0$

Proof of 4):

$$2a_2u^+ + \sigma - \mu w^+ = \sqrt{\sigma^2 + \frac{4a_2\alpha(\mu\phi\kappa + \chi\xi)}{a_1\xi}} - \frac{2\alpha\phi\kappa\mu a_2}{a_1\xi\left(\sqrt{\sigma^2 + \frac{4a_2\alpha(\mu\phi\kappa + \chi\xi)}{a_1\xi}} - \sigma\right)}$$

On simplifying to a fraction, denominator is positive.

Numerator:

$$a_1\xi\left(\sqrt{\sigma^2 + \frac{4a_2\alpha(\mu\phi\kappa + \chi\xi)}{a_1\xi}} - \sigma\right)\sqrt{\sigma^2 + \frac{4a_2\alpha(\mu\phi\kappa + \chi\xi)}{a_1\xi}} - 2\alpha\phi\kappa\mu a_2$$

$$= a_1\xi\left(\sigma^2 + \frac{4a_2\alpha(\mu\phi\kappa + \chi\xi)}{a_1\xi} - \sigma\sqrt{\sigma^2 + \frac{4a_2\alpha(\mu\phi\kappa + \chi\xi)}{a_1\xi}}\right) - 2\alpha\phi\kappa\mu a_2$$

$$= \left(a_1\xi\sigma^2 + 4a_2\alpha(\mu\phi\kappa + \chi\xi) - a_1\xi\sigma\sqrt{\sigma^2 + \frac{4a_2\alpha(\mu\phi\kappa + \chi\xi)}{a_1\xi}}\right) - 2\alpha\phi\kappa\mu a_2$$

$$= \left(a_1\xi\sigma^2 - a_1\xi\sigma\sqrt{\sigma^2 + \frac{4a_2\alpha(\mu\phi\kappa + \chi\xi)}{a_1\xi}}\right) + 4a_2\alpha\chi\xi + 2\alpha\phi\kappa\mu a_2$$

$$= a_1\xi\sigma\left(\sigma - \sqrt{\sigma^2 + \frac{4a_2\alpha(\mu\phi\kappa + \chi\xi)}{a_1\xi}}\right) + 2a_2\alpha(\mu\phi\kappa + 2\chi\xi)$$

$$= a_1\xi\sigma\left(\sigma - \sqrt{\sigma^2 + \frac{4a_2\alpha(\mu\phi\kappa + \chi\xi)}{a_1\xi}} + \frac{2a_2\alpha(\mu\phi\kappa + 2\chi\xi)}{a_1\xi\sigma}\right) > 0$$

Thus, numerator is non-negative.

Therefore $2a_2u^+ + \sigma - \mu w^+ > 0$ which implies,

$$2a_2u^+ + \xi u^+ + \sigma - \mu w^+ > 0$$

Therefore, the only sufficient condition for the local asymptotic stability of $(0,0,z^+,u^+,w^+)$ is:

$$\eta w^+ - \rho - a_2u^+ < 0$$

Proof of $\eta w^+ - \rho - a_2u^+ < 0$ if $\rho > \sigma$ and $\chi\xi + (\mu - \eta)\phi\kappa > 0$:

$$\eta w^+ - a_2u^+$$

$$= \frac{-1}{2}\left(-\sigma + \sqrt{\sigma^2 + \frac{4a_2\alpha(\mu\phi\kappa + \chi\xi)}{a_1\xi}}\right) + \frac{2\alpha\eta\phi\kappa a_2}{a_1\xi\left(-\sigma + \sqrt{\sigma^2 + \frac{4a_2\alpha(\mu\phi\kappa + \chi\xi)}{a_1\xi}}\right)}$$

$$= \frac{-a_1\xi\sigma^2 - 2\alpha a_2(\mu\phi\kappa + \chi\xi) + a_1\xi\sigma\sqrt{\sigma^2 + \frac{4a_2\alpha(\mu\phi\kappa + \chi\xi)}{a_1\xi}} + 2\alpha\eta\phi\kappa a_2}{a_1\xi\left(-\sigma + \sqrt{\sigma^2 + \frac{4a_2\alpha(\mu\phi\kappa + \chi\xi)}{a_1\xi}}\right)}$$

$$= \frac{-a_1\xi\sigma\left(\sigma - \sqrt{\sigma^2 + \frac{4a_2\alpha(\mu\phi\kappa + \chi\xi)}{a_1\xi}}\right) - 2\alpha a_2(\mu\phi\kappa + \chi\xi) + 2\alpha\eta\phi\kappa a_2}{a_1\xi\left(-\sigma + \sqrt{\sigma^2 + \frac{4a_2\alpha(\mu\phi\kappa + \chi\xi)}{a_1\xi}}\right)}$$

$$= \sigma - \frac{2\alpha a_2(\mu\phi\kappa + \chi\xi - \eta\phi\kappa)}{a_1\left(-\sigma + \sqrt{\sigma^2 + \frac{4a_2\alpha(\mu\phi\kappa + \chi\xi)}{a_1\xi}}\right)}$$

Thus, sufficient conditions for $\eta w^+ - \rho - a_2 u^+ < 0$ is $\rho > \sigma$ and $\chi\xi + (\mu - \eta)\phi\kappa > 0$.

Recall that $\rho = \frac{C_Y}{R_Y}\ln 2 + \frac{1}{p_Y}$ and $\sigma = \frac{C_U}{R_U}\ln 2 + \frac{1}{p_Y}$.

$$\rho > \sigma \Rightarrow \frac{C_Y}{R_Y}\ln 2 + \frac{1}{p_Y} > \frac{C_U}{R_U}\ln 2 + \frac{1}{p_Y}$$

$$\Rightarrow \frac{C_Y}{R_Y} > \frac{C_U}{R_U}$$

$$\Rightarrow C_Y R_U > C_U R_Y$$

Recall that $\mu = \frac{k_Y}{VR_U}\ln 2, \eta = \frac{k_Y}{VR_Y}\ln 2,\ \phi = k_X\Psi,\ \xi = \frac{k_Y}{V}, \kappa = (1 - k_Y h)$.

Therefore,

$$\chi\xi + (\mu - \eta)\phi\kappa > 0 \Rightarrow \frac{k_X\Psi(\ln 2)(1-k_Y h)k_Y}{V}\left(\frac{1}{R_U} - \frac{1}{R_Y}\right) + \chi\frac{k_Y}{V} > 0$$

Sufficient conditions for stability of $(0, 0, z^+, u^+, w^+)$, in other words for the invasion and subsequent fixation of the mutant phenotype in the resident-mutant system:

1) $\frac{C_Y}{R_Y} > \frac{C_U}{R_U}$
2) $\chi > \frac{k_X\Psi(\ln 2)(1-k_Y h)(R_U - R_Y)}{R_U R_Y}$

**Existence of $E_3(0, \tilde{y}, \tilde{z}, \tilde{u}, \tilde{w})$**

If the above-mentioned sufficient conditions hold, this rules out the existence of the rest point $(0, \tilde{y}, \tilde{z}, \tilde{u}, \tilde{w})$. If these sufficient conditions for stability of $(0, 0, z^+, u^+, w^+)$ are not satisfied, then $(0, \tilde{y}, \tilde{z}, \tilde{u}, \tilde{w})$ can also exist additional to $E_0$, $E_1$ and $E_2$ under some conditions.

The equilibrium point of the redefined system $E_3(0, \tilde{y}, \tilde{z}, \tilde{u}, \tilde{w})$ is defined as,

$$\tilde{y} = \Big(-{a_1}^{3/2}\xi^{3/2}\rho(\rho-\sigma)(\mu\rho-\eta\sigma) + a_1\xi(\rho-\sigma)(\mu\rho-\eta\sigma)\sqrt{a_1\xi\rho^2+4a_2\alpha\eta\kappa\phi} + a_2\alpha(\eta - \mu)\sqrt{a_1\xi\rho^2+4a_2\alpha\eta\kappa\phi}(-\eta\kappa\phi+\kappa\mu\phi+\xi\chi) + \sqrt{a_1}a_2\alpha\sqrt{\xi}(\eta^2\kappa\rho\phi - 2\eta\kappa\mu\rho\phi + \eta\xi(\rho-2\sigma)\chi + \mu\rho(\kappa\mu\phi+\xi\chi))\Big) / \big(2\sqrt{a_1}a_2\sqrt{\xi}(a_1\xi(\rho-\sigma)(\mu\rho-\eta\sigma) - a_2\alpha\kappa(\eta-\mu)^2\phi)\big)$$

$$\tilde{z} = \frac{\alpha}{a_1}$$

$$\tilde{u} = \frac{-\sqrt{a_1}\alpha\xi(\mu\rho+\eta(\rho-2\sigma))\chi + \alpha(-\eta+\mu)\sqrt{\xi}\sqrt{a_1\xi\rho^2+4a_2\alpha\eta\kappa\phi}\chi}{2\sqrt{a_1}(a_1\xi(\rho-\sigma)(\mu\rho-\eta\sigma) - a_2\alpha\kappa(\eta-\mu)^2\phi)}$$

$$\tilde{w} = \frac{\sqrt{\rho^2 + \frac{4a_2\alpha\eta\kappa\phi}{a_1\xi}} + \rho}{2\eta}$$

In this paper, we do not analytically study the stability of the rest point $(0, \tilde{y}, \tilde{z}, \tilde{u}, \tilde{w})$ due to its complexity and also because we are only concerned about the possibility of mutant invasion and the conditions required for the stability and fixation of such a mutant system with "consortium". However, we did some numerical investigation of all the possible equilibria.

**(SI 3) Numerical Analysis**

This section deals with the numerical simulations of the model to substantiate the analytical results. Let us first consider the following resident system:

$$\dot{x} = x(17.46 - 3x)$$

$$\dot{y} = y(0.023w - 3.106 - 2y)$$

$$\dot{w} = 80x - 0.01wy$$

The interior equilibrium point of the above system is $E_R(x^*, y^*, w^*) = (5.8, 22.4, 2076)$ when $\Psi = 400$, $a_1 = 3$, $a_2 = 2$, $k_X = 0.2$, $C_X = 4$, $R_X = 3$, $P_X = 10$, $k_Y = 0.1$, $R_Y = 0.3$, $C_Y = 1.2$, $P_Y = 3$, $\alpha = 17.46$, $\eta = 0.023$, $\rho = 3.106$, $\phi = 80$, $\xi = 0.01$, and $V = 10$.

*When $\rho > \sigma$ and $\chi = 30$ :*

Now we consider the coevolutionary system for the case when $\rho > \sigma$. For this let us define the following system:

$$\dot{x} = x(17.46 - 3x - 3z - 0.2u)$$

$$\dot{y} = y(0.023w - 3.106 - 2y - 2u)$$

$$\dot{z} = z(17.46 - 3x - 3z) + 0.2xu$$

$$\dot{u} = u(0.017w - 2.586 - 2y - 2u - 0.8x) + 30z$$

$$\dot{w} = 80x + 48z - 0.01wy - 0.01wu$$

Where $\beta = 0.8, h = 4, \chi = 30, R_U = 0.4, C_U = 1.3, \mu = 0.017, \kappa = 0.6, \sigma = 2.586$ and rest of the parameters are same as in the previously mentioned resident system.

The equilibrium points of the coevolutionary system are $E_0(0,0,0,0,0)$, $E_1(x^*, y^*, 0, 0, w^*) = (5.8, 22.4, 0, 0, 2076)$ and $E_2(0,0, z^+, u^+, w^+) = (0,\ 0,\ 5.8, 17.5, 1595)$

*When* $\rho < \sigma$ *and* $\chi = 5$ *:*

Now we consider the coevolutionary system for the case when $\rho < \sigma$. For this let us define the following system:

$$\dot{x} = x(17.46 - 3x - 3z - 0.2u)$$

$$\dot{y} = y(0.023w - 3.106 - 2y - 2u)$$

$$\dot{z} = z(17.46 - 3x - 3z) + 0.2xu$$

$$\dot{u} = u(0.022w - 3.149 - 2y - 2u - 0.8x) + 5z$$

$$\dot{w} = 80x + 48z - 0.01wy - 0.01wu$$

Where $\beta = 0.8, h = 4, \chi = 5, R_U = \ 0.32, C_U = 1.3, \mu = 0.022, \kappa = 0.6, \sigma = 3.149$ and rest of the parameters are same as in the previously mentioned resident system.

The equilibrium points of the coevolutionary system are:

$E_0(0,0,0,0,0)$, $E_1(x^*, y^*, 0, 0, w^*) = (5.8, 22.4, 0, 0, 2076)$, $E_2(0,0, z^+, u^+, w^+) = (0,\ 0,\ 5.8, 17, 1640)$ and $E_3(0, \tilde{y}, \tilde{z}, \tilde{u}, \tilde{w}) = (0, 5, 5.8, 12.2, 1623.7)$

**(SI 4) Probability of Evolutionary Substitution**

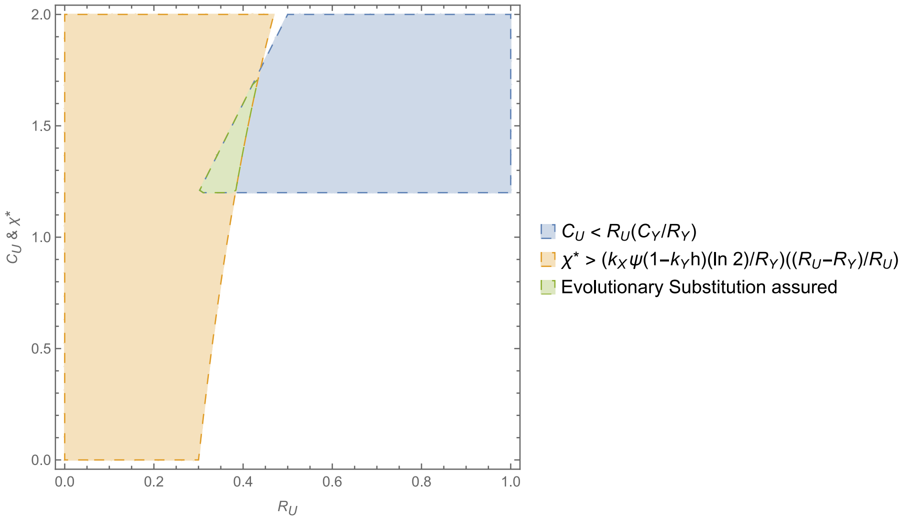


Consider (as in previous numerical analysis) $\Psi = 400$, $k_X = 0.2$, $k_Y = 0.1$, $R_Y = 0.3$, $C_Y = 1.2, h = 4$, and $\chi^*(\text{ Scaled } \chi\ ) = \chi/20$.

Area of intersection (green) = Area of intersection of $1.2 < y < 4x$ and $y > 5.55\left(\frac{x-0.3}{x}\right)$

$$= \int_{0.3}^{0.4387} 4x\, dx - \int_{0.3828}^{0.4387} 5.55\left(\frac{x-0.3}{x}\right) dx - 1.2(0.3828 - 0.3) = 0.022$$

Probability of evolutionary substitution ≥ 0.011.
